# “A mosquito bites and a butterfly flies": a specific response type of frontal patients in a similarity task

**DOI:** 10.1101/215020

**Authors:** Béatrice Garcin, Emmanuelle Volle, Aurélie Funkiewiez, Bruce L Miller, Bruno Dubois, Richard Levy

## Abstract

**Background:** Patients with neurodegenerative diseases affecting the frontal lobes have difficulties in categorization tasks, such as the similarity tasks. They give two types of unusual response to the question: “In what way are an orange and a banana alike?”, either a differentiation (“one is yellow, the other is orange”) or a concrete similarity (“they are sweet”).

**Objective:** To characterize the categorization deficit of frontal patients and develop a short diagnostic tool to assess the nature of these difficulties.

**Method:** We analyzed the responses provided by frontal and non-frontal neurodegenerative patients in a novel verbal similarity task (SimiCat). We included 40 frontal patients with behavioral variant fronto-temporal dementia (bvFTD) and progressive supranuclear palsy (PSP), 23 patients with Alzheimer’s disease (AD) and 41 healthy matched controls. Responses that did not correspond to the expected taxonomic category (*e.g.: fruits*) were considered as errors.

**Results:** All patients groups were impaired at the SimiCat test compared to controls. Differentiation errors were specific of frontal patients. Receiver operating characteristic analyses showed that a cut-off of two differentiation errors or more achieved 85% sensitivity of 100% specificity to discriminate bvFTD from AD. A short version of the test (<5 min) showed similar discriminative validity as the full version.

**Conclusion:** Differentiation responses were specific of frontal patients. The SimiCat demonstrates good discriminative validity to differentiate bvFTD and AD. The short version of the test is a promising diagnostic tool that will need validation in future studies.

## Introduction

Categorization represents a set of mental processes by which the brain classifies objects and events. The ability to categorize information has an impact on virtually all domains of cognition and behavior (Lawrence W. Barsalou, 1991). Categorization abilities can be assessed by the Similarities task (Dubois, Slachevsky, Litvan, & Pillon, 2000; Kaplan, 1991; Mattis, 1988; Wechsler, 2008), which is also referred to as a concept formation task. In this task, subjects have to categorize two concrete or abstract items (e.g.: “how are an orange and a banana alike?”) and give their taxonomic category (“fruits”).

Similarities task is often part of the clinical assessment of neurodegenerative patients. In particular, such a test is included in several batteries assessing executive functions (Dubois et al., 2000; Kramer & Quitania, 2007; Mattis, 1988; Wechsler, 2008). A deficit in the similarities task has been shown in the prodromal stages of neurodegenerative diseases (Fabrigoule et al., 1998) and correlates with measures of functional independence in dementia (Loewenstein, Rubert, Argüelles, & Duara, 1995), which makes it a useful assessment tool in dementia (Jacobs et al., 1995).

Patients with neurodegenerative diseases affecting the frontal lobes, notably the behavioral variant Fronto-Temporal Dementia (bvFTD) and Progressive Supranuclear Palsy (PSP)[10], show poor performances in Similarities task (Dubois et al., 2000; Kramer & Quitania, 2007; Lagarde et al., 2015). However, the reasons why frontal patients present with categorization difficulties are not well understood. Categorization is a complex neurocognitive function, relying on semantic knowledge and executive functions (T Giovannetti et al., 2001), including similarity detection in objects that are physically different, abstraction, and response selection according to the rule (Garcin, Volle, Dubois, & Levy, 2012). Each of these processes may contribute to the deficit and lead to distinct types of categorization problems. Consequently, the nature of the deficit may differ between patients with neurodegenerative diseases affecting the frontal lobes, such as bvFTD or PSP and patients with neurodegenerative diseases that affect more posterior regions, such as Alzheimers’disease (AD) (Tania Giovannetti et al., 2013).

Consistent with this hypothesis, clinical observations suggest that two types of incorrect responses are produced by frontal patients at the similarities task: a more concrete than expected similarity (e.g.:“they have a peel”) and a differentiation between the items (e.g.:“one is orange, the other is yellow”)(see video supplementary material of (Garcin et al., 2012)). These types of responses have been described in patients with a frontal neurodegenerative disease, and their measurement may help to discriminate frontal damage (bvFTD or PSP) from other neurodegenerative diseases Unfortunately, the typical scoring of the similarity task is quantitative: it only reports the severity of impairment, and fails to characterize the nature of the categorization deficit (Kaplan, 1991). A systematic description of the difficulties of patients with frontal neurodegenerative disease is necessary to precisely determine the nature of the deficit, and to stimulate further studies on the underlying mechanisms (Tania Giovannetti et al., 2013). The underlying mechanisms explaining patients’ errors are not known yet. One hypothesis is that differentiation responses in patients reflect difficulties in inhibiting a dominant mental representations (imagery) triggered by the items to be compared (Lagarde et al., 2015). Alternatively, patients may also have abstraction difficulties (Garcin et al., 2012)or difficulties to process abstract (less imageable) words.

The current study addresses these unexplored issues. The main objectives of our study were to characterize the categorization deficit of frontal patients (bvFTD and PSP) in terms of concrete similarities and differentiation errors, and to examine its specificity compared to non-frontal neurodegenerative patients (AD). The secondary objective was to develop a short diagnostic tool to target the specific difficulties of frontal patients. For these purposes, we developed a similarity tasks (the SimiCat) based on 28 pairs of taxonomically related words that we controlled for imageability, length and frequencies in French.

## METHOD

### Participants

We recruited patients from the multidisciplinary memory clinics of Saint-Antoine and Salpêtrière hospitals between November 2011 and July 2016. The patients met current diagnostic criteria for probable or definite bvFTD (Rascovsky et al., 2011), PSP (Litvan et al., 1996), or amnestic presentation AD (McKhann et al., 2011). They were not included if global testing showed severe impairment (i.e. Mini-Mental Status Examination (MMSE) score (Folstein, Folstein, & McHugh, 1975) < 16/30) or if neuropsychological testing revealed a significant semantic impairment according to the normative data of a French denomination task (Merck et al., 2011; Thuillard Colombo & Assal, 1992).

We recruited healthy controls from patients’ spouses in both memory clinics. They were matched to frontal patients (PSP and bvFTD) for age, sex and education. Exclusion criteria included history of mental illness, significant head injury, neurological conditions or substance abuse, and abnormal MMSE and/or Frontal Assessment Battery (FAB) score (Dubois et al., 2000).

The local ethics committee (CPP-IdF-Paris 5) approved the study and all the participants gave written informed consent.

### <u>Simi</u>larity-based <u>Ca</u>tegorization Task (SimiCat)

#### Rationale

The similarities subtest of the WAIS (Wechsler, 2008) is the most widely used similarities task. However, several issues limit its use in the clinic. First, it is based on a small number of items (n=19), which are heterogeneous in the nature of categorization processes. The items are linked in some cases according to taxonomic category (e.g. animals or fruits, n=9), but in others according to functional properties (e.g. food and fuel are two forms of energy); and/or general knowledge (e.g. honey and milk are produced by animals; silk and wool fiber originate from animals)(Wechsler, 2008). Second, the subtest has been designed to assess Intelligence Quotient (IQ) and correlates with education level and IQ (Longman, Saklofske, & Fung, 2007; Wechsler, 2008; Wisdom, Mignogna, & Collins, 2012). For this reason, there is a high variability of performance in healthy subjects (Harrison, Armstrong, Harrison, Lange, & Iverson, 2014; Wisdom et al., 2012), which is not appropriate for a clinical test that aims at distinguishing patients from healthy subjects with a clear difference required between both groups. Moreover, the subtest includes words of various imageability, sometimes intermingled in a same trial (such as poem and statue) that do not allow assessment of the impact of imageability on categorization abilities. For these reasons, we designed a new similarity task, named SimiCat.

### Material

The SimiCat task is based on 28 pairs of taxonomically related words (supplementary material 1). All words were controlled for length, and frequency (New, Pallier, Brysbaert, & Ferrand, 2004) (supplementary table 1). We selected pairs of words and categories that had high or low imageability, as measured by subjective ratings (Desrochers & Thompson, 2009). Words with low imageability such as “philosophy” are more abstract and may induce less mental imagery. There were three different kinds of pairs according to imageability of the pairs and categories (supplementary table 1). There were sixteen pairs of high imageability (HI) with a HI category linking them, 6 pairs of HI with a low imageability (LI) category linking them, and 6 pairs of LI with a LI category linking them.

#### Procedure

Instructions and items were given orally. Instruction was “what is the similarity between a ___ (e.g.: banana) and a ___(e.g.: orange)”. The participant’s responses were written down by the examiner. The participants received a feedback and correction only for the first item. When several answers were given, the participant had to select the one he considered his best answer. Instructions were repeated when no answer was given. We did not record response time.

#### Qualitative Analysis of the responses

The expected answer (correct answer) was the taxonomic category. Two other responses were analyzed: a more concrete than expected similarity (concrete similarity) and a differentiation between the items (differentiation). In order to classify each response, two examiners (BG and RL) analyzed the responses of 10 frontal patients, 10 AD patients, and 10 healthy controls and defined how to classify the responses. Then, they both classified all answers of all participants blindly to the participants’ condition. After a common definition of classification was determined, there was a high rate of consistency in the ratings (94.5%), and an agreement was found by discussion when rating differed between both examiners.

### Cognitive Assessment

General cognitive functioning was assessed using the MMSE (Folstein et al., 1975) and general executive functions were assessed by the FAB score (Dubois et al., 2000). Episodic memory was assessed by the 16 items free and cued recall test (Van der Linden et al., 2004). The patients also underwent a detailed neuropsychological testing that was part of the usual clinical assessment. The selection of tests varied according to the education level and nature of cognitive impairment, and systematically included an assessment of executive functions, visual episodic memory, praxis, and language.

### Statistical analyses

Data were analyzed using SPSS Statistics (V24.0) and significance was assumed at p<0.05. All variables were checked for normality of distribution using Kolmogorov-Smirnov tests. All variables but age followed a non-normal distribution. For this reason, we performed non-parametric analyses. Mann-Whitney U tests were used for paired-group comparisons and the Kruskal-Wallis test were used for comparison of more than two groups, followed when applicable by Mann-Whitney U tests for post-hoc pairwise comparisons with correction for multiple comparisons. Association between categorization scores and other measures were examined using Spearman’s correlation coefficients. Friedman tests were performed for repetitive measures analyses, followed by post-hoc Wilcoxon tests with correction for multiple comparisons if applicable. We also calculated receiver operating characteristic (ROC) curves to determine the best sensitivity and specificity indices of the task.

## RESULTS

### Demographic and clinical profiles

#### Comparison of frontal patients, AD patients and healthy controls (Table 1)

Frontal patients (n = 40: 20 bvFTD and 20 PSP patients), AD patients (n = 23) and healthy controls (n = 41) were matched for age, years of education, and gender. Frontal and AD patients were matched for disease duration. Both patient groups performed below controls in the general cognitive measure MMSE (all p values<0.001), but no difference was found between AD and frontal patients (p=0.66). AD and frontal patients also performed below controls on the general executive FAB score (all p values<0.001). In addition, frontal patients had lower FAB scores than AD patients (p<0.001). Finally, AD patients had significantly lower scores in the free and cued total recall score than frontal patients (p<0.001) (Table1).

**Table 1.** Comparison of Frontal patients, Alzheimer disease patients and healthy controls.

|  | Frontal patients | AD | Controls | Statistics |
| --- | --- | --- | --- | --- |
| Age (median (IQR)) | 71 (65-76.6) | 73.4 (67.9-76.6) | 69.4 (64.9-74.5) | KW (2)=1.2; p=0.33 |
| Education (median (IQR)) | 11.5 (9-15) | 14 (10.5-15) | 12 (9-15) | KW (2)=1.2; p=0.55 |
| Gender (% male) | 67.5% | 47.8% | 60.9% | Chi2=2.37; p=0.30 |
| Symptom duration<br>(median (IQR)) | 3.2 (1.6-5) | 4.2 (2.1-6) | NA | MW (U=388), p=0.31 |
| **MMSE (median (IQR))** | **25 (21.5-28)** | **24 (21.5-26)** | **29 (28-30)** | **KW (2)= 53.71 ; p<0.001** |
| **FAB (median (IQR))** | **12 (9-15)** | **15 (14-16)** | **17 (16-18)** | **KW (2)=54.57 ; p<0.001** |
| **Total recall (median**<br>**(IQR))** | **40 (33-45)** | **26 (9-31.5)** |  | **MW (U=111 ) ; p<0.001** |
| **SimiCat % correct**<br>**(median (IQR))** | **39.3 (20.5-67.9)** | **67.9 (60.7-**<br>**80.4)** | **92.9 (85.7-96.4)** | **KW(2)=65.7, p<0.001** |
| **% concrete similarities**<br>**(median (IQR))** | **17.9 (10.7-25)** | **17.9 (10.7-25)** | **3.6 (0-7.1)** | **KW(2)=37.5, p<0.001** |
| **% differentiation (median**<br>**(IQR))** | **21.4 (7.1-57.1)** | **0 (0-0)** | **0 (0-0)** | **KW(2)= 71.1 ,p<0.001** |
| **% other errors (median**<br>**(IQR))** | **7.1 (0-14.3)** | **7.1 (3.6-14.3)** | **3.6 (0-7.1)** | **KW(2)= 10.8, p=0.004** |

*IQR: inter quartile range; KW: Kruskal-wallis; MW: Mann-Whitney.*

### SimiCat performance

The first analysis of a subset of 10 frontal patients, 10 AD patients, and 10 healthy controls enabled us to identify six types of responses that we gathered into four categories: 1. <u>Correct response</u> was the expected taxonomic category (e.g.: “they are fruits”); the accuracy at the task was the percentage of correct responses; 2. <u>Concrete similarity:</u> the response designated either a characteristic shared by the items (feature similarity: “they are eatable”), or a contextual similarity (e.g.: “an orange goes well with a banana in a salad”); 3. <u>Differentiation</u>: the participant highlighted the distinctive characteristics of the objects to be compared instead of providing their similarity (e.g.: “the orange is orange and the banana is yellow”), sometimes in relation with the taxonomic category (e.g.: “they are different kinds of fruits”, 8% of differentiation errors); 4. <u>Other incorrect responses:</u> the response in this case was semantically wrong or no response was given (“I don’t know”). Concrete similarity, differentiation, and other wrong answers were all considered incorrect responses.

#### Between group comparison of accuracy (Table 1, figure 1)

Frontal patients were significantly impaired in comparison to AD patients (median accuracy in frontal: 39.3%, AD: 67.9%; U=176, p<0.0001), and AD patients were significantly impaired in comparison to controls (median accuracy in AD: 67.9%, controls: 92.9%; U=114.5, p<0.0001). There was a significant correlation between accuracy and education in controls (r=0.3, p=0.03), in frontal patients (r=0.44, p=0.005) but not in AD patients (r=0.04, p=0.8).

**Figure 1.**
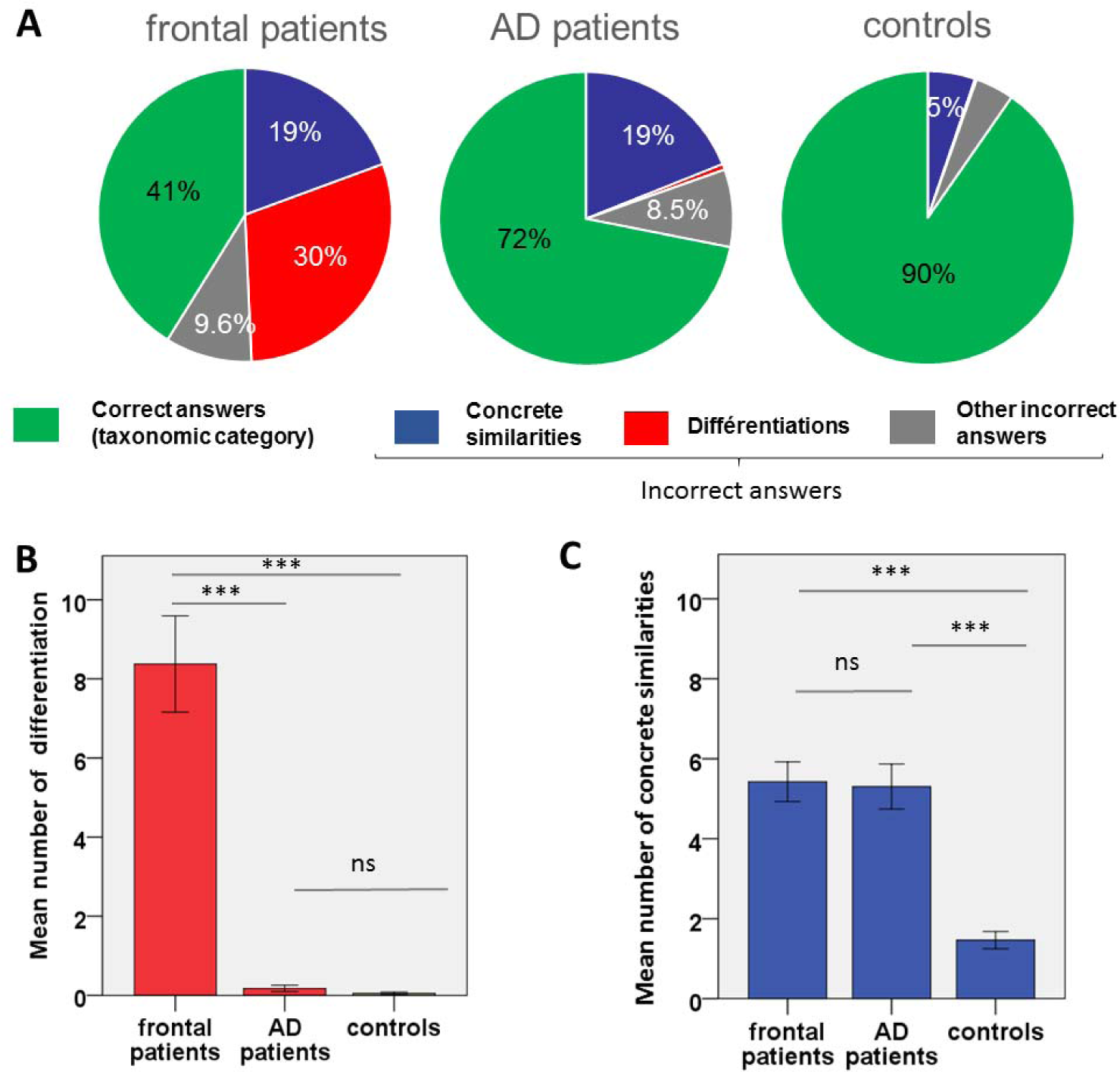
Repartition of responses in the participants’ groups. A. Repartition of all kinds of responses in the participants’ groups in mean percentages. B. Comparison of the number of differentiation errors in the groups of participants: mean +/-standard error of the mean (out of 28 pairs of words). Kruskal-Wallis test was significant (KW=71.1, p<0.000), Mann-Whitney U tests were used for paired comparisons. C. Comparison of the number of concrete similarities in the groups of participants: mean +/-standard error of the mean (out of 28 pairs of words). Kruskal-Wallis test was significant (KW=37.5, p<0.000), Mann-Whitney U tests were used for paired comparisons. ns: not significant; ***:p<0.001

#### Between-group analysis of errors (Table 1 and 2, and figure 1 and 2)

##### Differentiations

Differentiation responses were common in frontal patients (mean: 30%, median: 21.4%), but were almost never observed in controls or in AD patients (frontal vs. AD: U=50.5; P<0.001, frontal vs. controls: U=69.5 p<0.001). The proportion of differentiation responses did not significantly differ between bvFTD and PSP patients (U=157, p=0.3; table 3 and sup fig 1), nor did it differ between AD patients and controls (U=412.5, p=0.1). The number of differentiation errors correlated negatively with the severity of the frontal syndrome in the frontal group (r= −0.566, p<0.000), but not in the AD group (r=-0.125, p=0.57; only 4 values available for this analysis). Frontal patients and AD patients were split into three groups of increasing severity according to the FAB score (<8: very severe, 8-11: intermediate, and 12-15: mild). The similarity subscore of the FAB was removed from the total FAB score to avoid repetition with the SimiCat test. This allowed us to compare 17 frontal patients to 16 AD patients of similar mild frontal syndrome severity. In this subgroup, frontal patients gave significantly more differentiation responses than in the AD group, although FAB scores and MMSE scores were not significantly different between the groups (Figure 2 and Table 2).

**Table 2.**
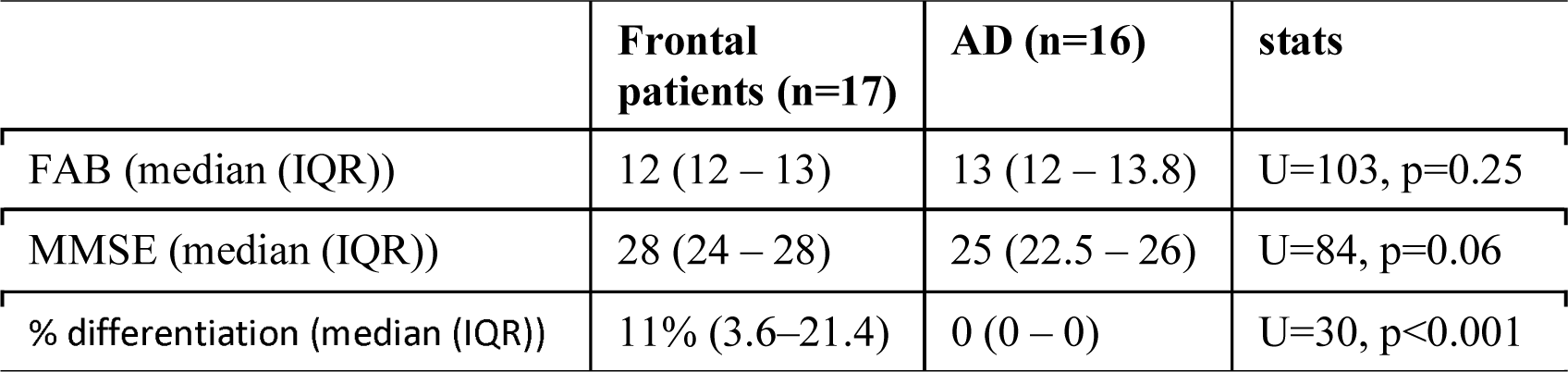
Comparison of Alzheimer and frontal patient subgroups with a mild frontal syndrome.

**Table 3.** Comparison of PSP and bvFTD patients.

|  | PSP | bvFTD | Statistics |
| --- | --- | --- | --- |
| **Age (median (IQR))** | **73.37 (68.6-78.8)** | **65.4 (63.5-72.3)** | **MW (U=106); p=0.01** |
| Education (years) (median (IQR)) | 10 (7 -15) | 12 (9 -17); | MW (U=181); p=0.6 |
| Gender (% male) | 65% | 70% | Chi2=0.73; p=1 |
| **Symptom duration (median; IQR)** | **2.6 (1.05-4.6)** | **3.5 (2.6-5.7)** | **MW (U=127); p=0.048** |
| MMSE (median; IQR) | 25 (24-28) | 25 (21-28) | MW (U=178.5); p=0.56 |
| FAB (median; IQR) | 12 (9-15) | 13 (8-15) | MW (U=181.5); p=0.62 |
| Total recall* (median; IQR) | 40 (35-47) | 40 (32-45) | MW: (U=124.5); p=0.54 |
| SimiCat % correct (median (IQR)) | 46 (18.7-67.9) | 37.5 (13.4-70.5) | MW (U=182); p=0.64 |
| % concrete similarities (median (IQR)) | 17.9 (11.6-25) | 14.3 (10.7-21.4) | MW (U=169); p=0.41 |
| % differentiation (median (IQR)) | 14.3 (4.5-38.4) | 7.5 (10.7-59.8) | MW (U=157); p=0.25 |
| % other errors (median (IQR)) | 10.7 (3.6-14.3) | 3.6 (0-9.8) | MW (U=182); p=0.64 |

*IQR; Inter-quartile range, PSP: Progressive Supra-nuclear Palsy, bvFTD : behaviour variant Fronto-temporal Dementia, MW: Mann Whitney.*

**Figure 2.**
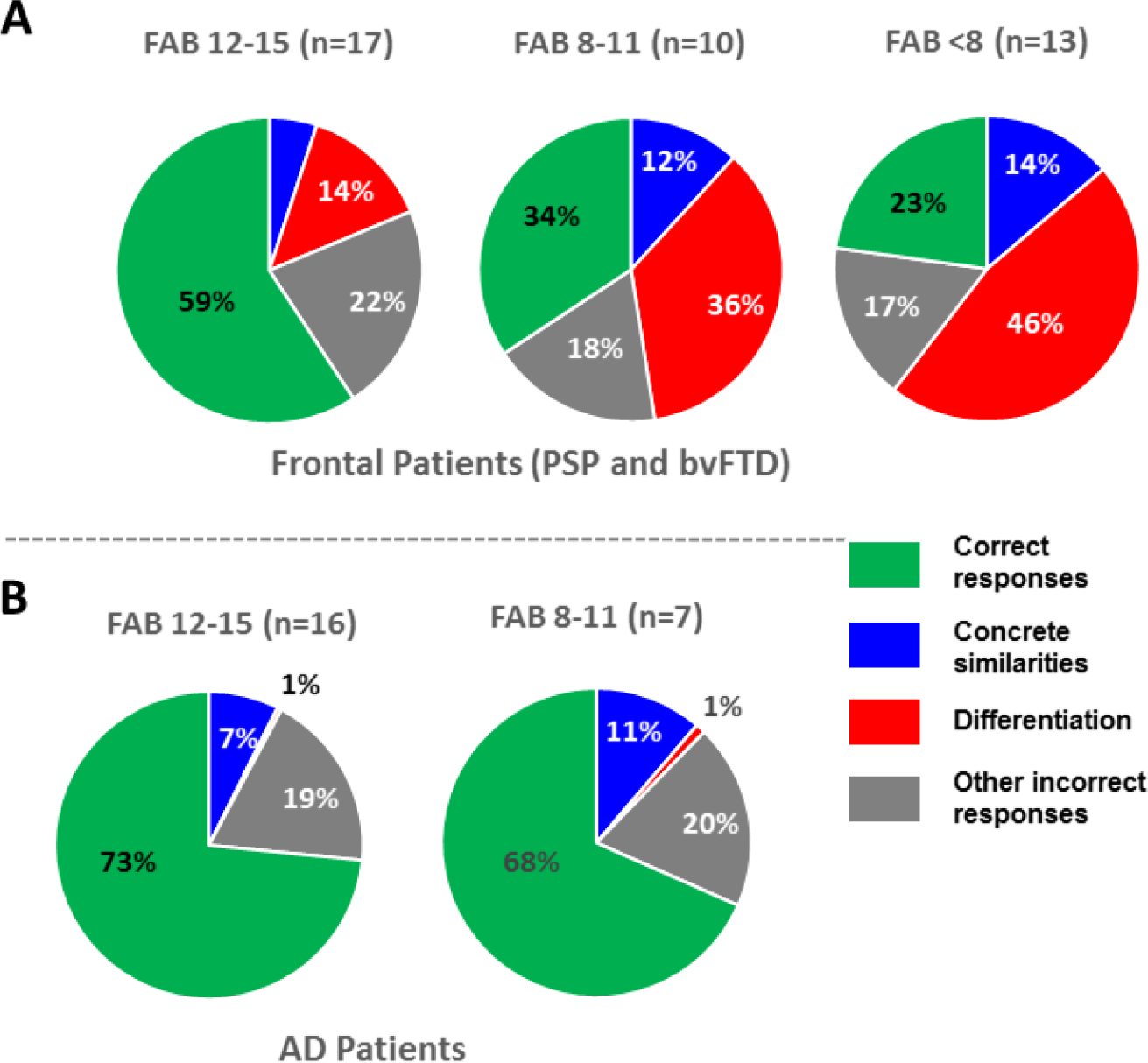
Repartition of responses according to severity of the frontal syndrome on the FAB score. Three groups of increasing severity were formed according to the FAB score ranging from 0 (very severe frontal syndrome) to 15 (no frontal syndrome). The similarity subscore was not included because repetitive with the Simicat test. Cut-off FAB scores were proposed to balance the proportion of patients in each group (according the 33th and 66^th^ percentiles of the FAB score in the group of patients). A. Repartition of answers according to the severity of the frontal syndrome in frontal patients. B. Repartition of answers according to the severity of the frontal syndrome in Alzheimer disease patients.

##### Concrete similarities

Concrete similarity responses were more common in both patient groups (mean: 19%, median: 17.9% in both groups) than in controls (mean: 5%, median 3.6%) (p<0.001 for both comparisons). AD and frontal patients did not differ in this score (p=0.98), nor did bvFTD patients and PSP patients (p=0.4 Table 3 and sup fig. 1).

##### Other types of incorrect responses

Other errors were more common in AD and frontal patients than in controls (p=0.004 and p=0.012 respectively), but there was no difference between AD and frontal patients (p=0.72).

#### Comparison of PSP and bvFTD patients (Table 3)

The proportion of correct responses and the proportion of each different kind of incorrect responses did not significantly differ between bvFTD and PSP patients (Table 3; Sup fig. 1).

### Performance of frontal patients according to imageability of the items (sup fig 2)

Frontal patients gave significantly more correct answers to high imageability (HI) pairs of words linked by a high imageability category, than in both low imageability (LI) category conditions (p<0.001 for both comparisons). There gave more differentiation errors to the HI pairs of words linked by an LI category in comparison to the HI/HI condition (p=0.002). Finally, more other errors were found when LI words were linked by an LI category (p=0.001, and p<0.001).

#### SimiCat differentiates bvFTD from AD

Differentiation errors were common in frontal patients and were not observed in AD patients, suggesting that this measure could be used to discriminate bvFTD from AD patients. We tested whether the number of differentiation responses can be used as a diagnostic tool to discriminate bvFTD from AD patients. The ROC analysis using the total number of differentiation responses revealed an area under the curve of 0.937 (95% CI: 0.851-1; p=0.000) with a sensitivity of 85% and specificity of 100% at a cut-off score of 2 or more.

#### Development of a short version of SimiCat (Table 4, supplementary material)

With the aim to develop a short SimiCat version for clinical use, we selected ten pairs and tested the accuracy of this short test to discriminate bvFTD from AD. We selected six pairs for which differentiation responses were 100% specific to frontal patients. For those six pairs, 2 points were attributed to each differentiation answer. We additionally selected four pairs of the original task that were highly sensitive to differentiate frontal from AD patients, but less specific. 1 point was attributed for a differentiation error to those pairs. This led to a “differentiation score” rated on 16 points for the short task of 10 items. The ROC curve analysis of the differentiation score revealed an area under the curve of 0.937 (95% CI: 0.852-1; p=0.000). Cut-off scores derived from this analysis indicated that a total score of 1 or more differentiation errors identified bvFTD with 90% sensitivity and 87% specificity, while a total score of 2 or more achieved 80% sensitivity and 100% specificity. A codebook with all classified participants’ responses at the SimiCat-10 is provided as supplementary material 2.

**Table 4.** Short version of SimiCat test for clinical use to discriminate AD from bvFTD.

| Pairs of words | Points for a differentiation error |
| --- | --- |
| Horse - turtle | 2 |
| Pizza-chocolate | 2 |
| Boat-car | 2 |
| Tobacco-alcohol | 2 |
| Surgery-Medication | 2 |
| One century- 3 seconds | 2 |
| Sailor-physician | 1 |
| March-June | 1 |
| Jealousy-friendship | 1 |
| history-philosophy | 1 |

## Discussion

In this study, we used a new Similarities task named “SimiCat” to analyze the nature of categorization deficits of neurodegenerative patients. We found two main kinds of errors in frontal patients: differentiation and concrete similarity. Only bvFTD and PSP patients produced differentiation errors, while all groups of participants made concrete similarities errors, suggesting that separate mechanisms, relying on distinct brain circuits, may underlie both error types. We propose a short version of the test “SimiCat-10” as a diagnostic tool to differentiate between AD and bvFTD with excellent discriminative validity.

<u>**The first new finding** of the current study is that differentiation responses</u> were specific to frontal patients. When providing differentiation responses, patients did not follow the instruction (that is to give the most relevant similarity between the items), and instead described the items differences or the items most prominent characteristic. Several cognitive dysfunctions may explain this behavior: *First, frontal patients may have an inability to follow and maintain the rule*. Using a spatial planning task, Carey et al. ((Carey et al., 2008)) showed that frontal patients had an increased propensity for rule violation in comparison to AD patients although their overall performances were similar. The failure to adhere to rules may stem from a number of different reasons including inattentiveness, impulsivity, disinhibition, that are common symptoms in bvFTD (Rascovsky et al., 2011). *Second, they may have a specific deficit in similarity detection,* defined as the ability to perceive the common features of two items (Garcin et al., 2012) despite their differences. *Finally, they may have difficulty in the inhibition of dominant mental representations (imagery) triggered by the items to be compared* (L. W. Barsalou, 1999). In agreement with this hypothesis, it is noteworthy that frontal patients produced more differentiation errors for the HI/LI condition. In this condition, the items to be compared were of high imageability and likely to induce high mental imagery, while the category they had to find was of low imageability and less accessible to mental imagery. In other words, the HI/LI condition required more inhibition of strongly induced but irrelevant mental images of the concrete items to be compared.

Rule representation/maintenance, similarity detection and inhibition of mental representations are all part of the executive functions, which rely on the lateral PFC’s integrity (Stuss & Alexander, 2000). In this way, differentiation responses can be considered a consequence of a general dysexecutive syndrome, due to lesions of the PFC and/or its connections. However, although to a lesser extent, AD also affects frontal lobes (Migliaccio et al., 2015), and induces a dysexecutive syndrome (Perry & Hodges, 1999). In subgroups sharing similar dysexecutive syndrome severity, frontal patients provided significantly more differentiation errors than did AD patients. For these reasons, we believe that differentiation responses are not only the consequence of a general dysexecutive disorder, but represent a specific behavior of bvFTD and PSP patients, due to the regional specificity of the neurodegenerative lesions in these diseases (Lagarde et al., 2013; Rosen et al., 2002). Similarity detection, rule representation/maintenance and inhibition are thought to rely on the ventrolateral PFC (Bengtsson, Haynes, Sakai, Buckley, & Passingham, 2009; Bunge, 2004; Garcin et al., 2012; Hampshire, Chamberlain, Monti, Duncan, & Owen, 2010). Further studies will be necessary to precise the mechanisms and brain networks responsible for differentiation responses, and to determine the role of the ventrolateral PFC (inferior frontal gyrus).

By contrast to differentiation responses, <u>**concrete similarities** errors were seen in all participants groups</u> with a higher proportion of these responses in patients than in healthy controls. The proportion of concrete similarity responses was similar in both patient groups (AD and frontal patients), and was therefore nonspecific as to the neurodegenerative disease. Various mechanisms may explain these responses: deficit in abstract thinking abilities (Garcin et al., 2012), impaired semantic knowledge (T Giovannetti et al., 2001) and deficits in response selection (T Giovannetti et al., 2001). The mechanisms explaining concrete similarities errors may differ in frontal patients and AD, and their neural substrate have to be determined.

### Comparison of PSP & bvFTD

Based on recent studies ((Brenneis et al., 2004; Lagarde et al., 2013)) showing high similarities in behavioral and atrophy patterns in PSP and bvFTD, we decided to pool patients suffering from both diseases in the frontal group. Comparison of PSP and bvFTD showed no significant differences in accuracy on the SimiCat test and in general neuropsychological tests, confirming a behavioral similarity between groups. Moreover, there was no difference in the proportion of differentiation responses in PSP and bvFTD, suggesting a similar mechanism by which categorization was altered in both groups, and a damage to similar brain networks supporting this mechanism.

### Clinical implications: SimiCat-10: a new screening tool?

**The second new finding** of this study is that the SimiCat test discriminates frontal and non-frontal neurodegenerative patients with a high degree of accuracy. Neuropsychological differentiation between bvFTD and AD remains challenging in clinical settings given the widespread overlap of cognitive profiles (Hutchinson & Mathias, 2007; Ritter, Leger, Miller, & Banks, 2016). Accurate diagnosis is important because of the implication for prognosis, heritability and therapeutic interventions. Several tests have been recently developed to address this issue such as the Social cognition and Emotional Assessment Battery (Funkiewiez, Bertoux, de Souza, Lévy, & Dubois, 2012) or the FRONTIER Executive Battery (Leslie et al., 2015). In this line, the SimiCat test has a high discriminative validity to differentiate bvFTD and AD. We propose a short version of this test named SimiCat-10. Compared to other tests, the Simicat-10 is short and easy to use in the clinic: it takes less than 5 minutes, and does not require any specific equipment. Future studies will be necessary for validation of the short version. They may explore the combination of SimiCat-10 with social cognition measures and executive tests for accurate bvFTD diagnosis.

This study has several limitations. First it was performed in neurodegenerative disease and it remains unknown whether non-degenerative frontal pathologies such as traumatic brain injury, strokes or brain tumors have similar impact on categorization abilities as frontal neurodegenerative diseases. The SimiCat test would be a good tool to assess these disorders, and to help adapt the cognitive rehabilitation programs. Second, the validity of the SimiCat-10 for differentiating bvFTD and AD was assessed on the same patient population as the full version, and future studies within large patient populations are needed to directly evaluate the differential diagnostic properties of the SimiCat-10 to differentiate dementia subtypes. Finally, this test was developed in French, and validation of its discriminative accuracy will be necessary in other languages, notably in English.

In summary, we showed that differentiation errors are specific to frontal patients. The Short version of the test (SimiCat-10) is a promising and easy test to differentiate between bvFTD and AD.

## Acknowedgements

The authors would like to thank the participants and their families for participating in our research. We thank Nicolas Defoor, Valentine Facque, and Céline Chamayou for their help in the inclusion of patients.

## Contributors

BG, EV and RL contributed to the design and conceptualization of the study, analysis, and interpretation of data, drafting and revising the manuscript. BG and AF contributed to data acquisition. BD and BM contributed to interpretation of data and revision of manuscript.

